# The BMP4-BMPR1A axis represses BRCA1 in mammary stem cells and contributes to tumor initiation

**DOI:** 10.1101/2025.06.27.661450

**Authors:** Simon Aho, Marie Perbet, Isabelle Treilleux, Adrien Buisson, Sandrine Jeanpierre, Emmanuel Delay, Véronique Maguer-Satta, Boris Guyot

## Abstract

Basal-like breast cancer (BLBC) is an aggressive subtype frequently characterized by homologous recombination deficiency (HRD) and BRCAness, even in the absence of BRCA1 mutations. Here, we identify the BMP4-BMPR1A signaling axis as a novel regulator of BRCA1 expression and a driver of BRCAness in non-transformed immature mammary cells. We observed that high BMPR1A expression is correlated with low BRCA1 levels and poor BLBC prognosis in patients. We show that exposure to BMP4 induces BRCA1 transcriptional repression via BMPR1A, promoting a basal differentiation and impairing homologous recombination. This results in increased sensitivity to PARP inhibitors (PARPi) and accumulation of genetically-unstable, immature cells. In addition, long-term BMP4 stimulation or BMPR1A overexpression induced transformation. Our findings uncover a mechanism by which the tumor microenvironment may contribute to BLBC initiation through BMP4-induced suppression of BRCA1, and highlight BMP signaling and the resulting BRCAness as a therapeutic vulnerability in wild-type BRCA1 BLBCs.

## Introduction

Breast cancer is a complex disease encompassing several molecular subtypes defined by distinct gene expression profiles. The molecular classification distinguishes four intrinsic subtypes, namely luminal A, luminal B, normal-like, HER2 enriched and basal-like (BLBC)^1–3^. BLBCs represent 20% of all breast cancer cases^3–5^ and display the worst prognosis^6–9^. They express genes characteristic of myoepithelial cells, progenitor and mammary stem cells from the basal layer of the mammary gland^10^, as well as genes related to epithelial-mesenchymal transition (EMT), proliferation and stemness^1,3^. Interestingly, BLBC are enriched in cancer stem cells (CSC) with a CD44+/CD24-phenotype^11^. CSC have similar self-renewal and differentiation properties as healthy mammary stem cells (MaSC), but also exhibit tumorigenic and treatment-resistant features. They have thus been implicated in early stages, as well as in the progression and relapse of a number of cancers, including breast cancer^12^. Though the cell of origin of BLBC remains controversial, MaSC and an ER^-^ luminal progenitors have been proposed^13,14^.

The histological classification of BLBC resembles that of the highly aggressive triple-negative breast cancers (TNBC) ^6,15^, as they display a low level of expression of estrogen and progesterone receptors (ERα and PR), and no HER2 enrichment (human epidermal growth factor receptor-2)^1,3^. BLBCs are also positive for basal cytokeratins (CK5/6, CK14, CK17) and EGFR^16,17^. BLBCs frequently exhibit specific mutational patterns and chromosomal alterations, resulting from a defective homologous recombination (HR)^18^; the most error-free DNA double-strand break repair pathway essential for maintaining genome stability^19^. Among the genes involved in HR, the best known are BRCA1 and BRCA2, the germline mutations of which are strong risk factors for ovarian and breast cancers^20^. Aside from these mutations, alternative mechanisms of homologous recombination deficiency (HRD) can result in a “BRCAness” phenotype (*i.e.*, a phenocopy of these inherited mutations^21^). Interestingly up to 80% of BLBCs present an HRD^18,22–24^, stemming from germline mutations in BRCA1 (10%) or BRCA2 (2.5%), somatic mutations in BRCA1 (6%) or BRCA2 (2.5%) ^25^, hypermethylation of the BRCA1 promoter (14^26^-34%^27^) or overexpression of ID4 (37.5-50%)^28^. In addition, miRNAs targeting the BRCA1 mRNA have also been described^29–33^.

At the level of the tumor microenvironment (TME), it was shown that some cytokines, including TGFβ, could promote HRD in tumors^34,35^, arguing in favor of their implication in driving BLBC carcinogenesis. Bone morphogenetic proteins (BMP) are part of the TGFβ superfamily of cytokines and are secreted by the mammary gland microenvironment. BMPs activate transmembrane serine-threonine kinase receptors (BMPR)^36^, which in turn induce SMAD1/5/8 phosphorylation and binding to SMAD4. The complex then translocates to the nucleus and regulates target gene expression. SMAD-independent pathways involving the activation of MAP kinases, such as ERK, JNK or p38^37^, have also been reported.

Although BMP pathway alterations have been reported in luminal breast cancers tumorigenesis^38^, their involvement in BLBC carcinogenesis is limited to a few studies suggesting a role for BMPR1A and/or BMP4 ^39,40^. Here we demonstrate that the BMP4/BMPR1A signaling axis decreases BRCA1 gene expression, resulting in an increased basal differentiation of mammary progenitors and in HRD associated with genomic instability and transformation.

## Methods

### Cell culture

The MCF10A cell line (RRID:CVCL_0598) was supplied by ATCC (the American Type Culture Collection, USA) and grown in Dulbecco’s Modified Eagle’s Medium (DMEM)/F-12 with Penicillin/Streptomycin (Life Technologies), 1% GlutaMax (Thermo Fisher), 5% Horse Serum (Life Technologies), 10 µg/mL insulin (Novorapid), 0.5 µg/mL Hydrocortisone (Sigma), 100 ng/mL Cholera Toxin (Sigma) and 20 ng/mL Epithelial Growth Factor (EGF) (Sigma) in 5% CO2 at 37°C. Cell cultures were routinely tested for mycoplasma contamination. BMP4 was used in the 10-50 ng/mL range as indicated. The LDN-193189 BMP receptor inhibitor (Sigma-Aldrich) was used at 100 ng/mL and the E6201 inhibitor (Spirita Oncology, LLC) at 100 nM. E6201 is largely used as a MEK1 and FLT3 inhibitor but is also a preferential BMPR1A inhibitor (IC₅₀ = 0.45 nM for BMPR1A; 36 nM for BMPR1B) (unpublished supplier data). To estimate the sensitivity of Olaparib and Talazoparib, the two PARPi (Sigma Aldrich) were used at the 0.5-4 µM and 1-5 nM ranges, respectively.

### Generation of cell lines

For MCF10A cells with inducible shRNA-mediated BRCA1 knock-down and their control, the LT3GEPIR-shLuc construct (kind gift of Dr Johannes Zuber) was used and the shBRCA1.661^41^ sequence was inserted in place of the shLuciferase sequence. Cells were transfected using a Neon (Invitrogen) system with the LT3GEPIR-shBRCA1.661 or the LT3GEPIR-shLuc as a control. 3 days after transfection, cells with a stable integration were selected with 0.5 µg/mL puromycin for 4 days. Resistant cells were treated overnight with 400 ng/ml doxycycline. GFP-positive cells were FACS-sorted and further expanded without doxycycline. For BMPR1A over-expression, MCF10A were transfected using a Neon apparatus with a BMPR1A expression vector (pcDNA3-HA-BMPRIA WT) (kind gift from Dr. Peter ten Dijke) or an empty control vector (pcDNA3-HA) and selected with puromycin (0.5 µg/mL). For MCF10A-FUCCI cells, the mVenus-hGeminin/pcDNA3 and mCherry-hCdt1/pcDNA3 plasmids from the Riken Institute were transfected into MCF10A cell line before selection with puromycin (1 µg/mL).

### Flow cytometry and Fluorescence-activated cell sorting

Flow cytometry was performed on an LSR Fortessa cytometer. The Diva software was used for acquisition and the Flow-Jo software for analysis. Cells sorting was performed using a BioRad S3e cell sorter (RRID:SCR_019710).

### BMP4 ELISA in primary tissues

Tumor samples and normal tissue supernatants were obtained as described^36^ and soluble BMP4 quantification was performed by ELISA following the manufacturer’s instructions (Quantikine ELISA Kit; R&D Systems).

### Functional assays

For epithelial colony forming cell (E-CFC) assay, 250 cells of interest were added to 8.10^4^ 30 Gy-irradiated NIH3T3 (RRID: CVCL_0594) feeder cells in 12-well plates in MCF10A medium. After 6 days of co-culture, colonies were fixed with ice-cold methanol and stained using Wright’s stain (Sigma-Aldrich). The different colony types were then counted under a 4x magnification according to their ultrastructural features^42^. For mammosphere assay, cells were grown in 96-well ultra-low attachment plates using mammary epithelial cell basal growth medium (MEBM) with B27 additive (Thermo-Fisher), 20 ng/mL bFGF (Life Technologies) and 4 mg/mL heparin (Life Technologies). After 6 days, mammospheres were counted at 4x magnification. For soft-agar colony formation assay 1 mL of MCF10A culture medium containing 0.75% of agar was poured into 12-well culture plates and incubated at room temperature for 1 h. 1 mL of MCF10A culture medium containing 0.45% of agar and 45,000 cells was then poured on top. After incubation at room temperature for 30 min, cells were kept at 5% CO2 and 37°C for 1 to 2 weeks and counted under a Zeiss microscope at 4x magnification.

### Immunohistochemistry

IHC analyses were conducted on formalin-fixed triple-negative breast cancer samples at diagnosis. H&E and IHC staining of paraffin sections was carried out using standard methods and anti-BMPR1A (MA5-17036, Invitrogen; RRID:AB_2538508) and anti-cytokeratin 14 (Clone SP53, 06732429001, Roche diagnostics) antibodies. The percentage of positive cells and the staining intensity were measured and an expression score defined as the product of these two values.

### Immunofluorescence

Cells (1500 per well) were seeded in 96-wells Phenoplates (Revvity) with the indicated treatments and grown for 48 hours. DNA DSB were generated with a 2Gy dose using a X-ray irradiator. At various time-points post-irradiation, cells were fixed with 4% formaldehyde for 15 minutes at room temperature (RT) before permeabilization for 3 minutes at RT with 1% Triton-X100 in PBS. After a 1 hour blocking step in PBS 5% BSA at RT, cells were incubated with the primary antibody for 2 hours at RT in PBS 5%BSA, washed three times in PBS, incubated with the secondary antibody in the same buffer for 1 hour at RT with Hoechst 33342 and washed again. Labeled cells were imaged using an automated confocal microscope (Opera Phenix Plus, Revvity, RRID:SCR_021100) with a 40x magnification. Images obtained after maximum projection of the z-stacks were analyzed using the Harmony software (RRID:SCR_023543) to determine the number of 53BP1, RAD51 and γH2AX foci per nucleus.

Mouse anti-53BP1 (Millipore MAB3802, RRID:AB_11212586, 1/500)

Rabbit anti-RAD51 (Abcam ab133534, RRID:AB_2722613, 1/400)

Rabbit anti-γH2AX (Cell Signaling 5438, RRID:AB_10707494, 1/1000)

Goat anti-rabbit IgG-Alexa Fluor 647 (Invitrogen A21245, RRID:AB_2535813, 1/1000)

Rabbit anti-mouse IgG-Alexa Fluor 555 (Abcam ab150126, 1/1000)

### Reverse transcription and quantitative polymerase chain reaction

Total RNA was extracted using a RNeasy Mini Kit (Qiagen). Quantity and quality of RNA were measured using a spectrophotometry. cDNA synthesis was performed on 1 μg of RNA using the SuperScript™ RT reagent kit (Thermo Fisher Scientific). qPCR was performed using the QuantiFast SYBR® Green RT-PCR Kit (Qiagen) on a LightCycler480 (Roche Diagnostics, RRID:SCR_018626) (95°C for 5 min, 45 cycles of 95°C for 10 s and 60°C for 30 s). Negative controls (no template or no reverse transcriptase) were included. The expression of the genes of interest were normalized against the CPB and ACTB1 genes expression using the ΔΔCT method.

### Western-blot

One million cells were lysed in 100 μl of Laemmli buffer. Samples were boiled for 5 min at 100°C and sonicated for 10 seconds. 30 μl of lysate were run on 6% SDS-PAGE in 25 mM Tris, 192 mM glycine, 0,1% SDS and transferred to PVDF membranes. Membranes were saturated in 5% BSA in TBS-T (1% Tween-20) (Euromedex). Primary antibodies were incubated overnight at 4°C and washed with TBS-T. Incubation with HRP-conjugated secondary antibodies was performed at room temperature for 1 h before washes with TBS-T buffer. Clarity™ Western ECL Substrate (Biorad) and Chemidoc™ Imaging System (Biorad, RRID:SCR_019037) were used to visualize the antibodies. Signals were quantified using the ImageLab software (RRID:SCR_014210).

Mouse anti-BRCA1 (Merck OP92, clone MS110, 1/500)

Rabbit anti-cleaved Caspase 3 (Cell signaling 9661, RRID:AB_2341188, 1/1000)

Mouse anti-GAPDH (Abcam Ab8245, RRID:AB_2107448, 1/10000)

Rabbit anti-β-tubulin (Abcam, Ab6046, RRID:AB_2210370, 1/2000)

### Comet assay

Comet assays were performed using the CometAssay Single Cell Gel Electrophoresis Assay kit (Trevigen). Cells were resuspended in PBS at 1.5×10^6^/mL. 12.5 µL of the cell suspension was diluted in 125 µL of 1% low-melting point Agarose. 40 µL of cell suspension was spread on a microscope slide at 37°C before incubation for 30 min at 4°C in a dark, humid chamber. The slides were then incubated in a lysis solution at 4°C for 1 h. For the neutral comet assay, slides were incubated in 1x neutral electrophoresis buffer at 4°C for 30 min and run in 1x neutral electrophoresis buffer for 45 min migration at 21 volts and 4°C. After incubation in a precipitation solution for 30 min at room temperature, the slides were incubated for 30 min in 70% ethanol and dried. For the alkaline comet assay, slides were placed in alkaline buffer at 4°C for 40 min and run in 1x alkaline buffer for 30 min at 25 volts and 4°C. The slides were then neutralized for 3×10 min at 4°C, fixed in 70% ethanol for 30 min and dried. After DNA staining in a SyBr Gold solution at room temperature for 30 min, slides were mounted with Fluoromount medium (RRID:SCR_015961). Nuclei were imaged using a fluorescence microscope (Zeiss Axio Imager M2, RRID:SCR_024706) and analyzed on ImageJ (RRID:SCR_003070) using the opencomet (RRID:SCR_021826) program.

### In silico analysis

Evaluation of BMPR1A, BMPR1B, BMP2 and BMP4 mRNA levels in breast cancer molecular subtypes and healthy tissue controls were done using the Gene Expression-Based Outcome for Breast Cancer Online (http://co.bmc.lu.se/gobo) and the Breast Cancer Gene-Expression Miner v5.0 (bc-GenExMiner v5.0) tools. Correlation analyses were performed using bc-GenExMiner v5.0 and Kaplan-Meier plots for survival analyses were performed with the kmplotter (https://kmplot.com/analysis/) tool.

### Statistical analysis

The relationships between BRCA1 transcriptional and BMPR1A protein expression was modeled using linear regression. The influence of confounding factors (CK14 positivity, age at diagnosis and stage) was evaluated with bivariate regression analysis. Stata software (version 18) (Stata 2023) and R (R Core Team 2023) were used for the analyses. For PARPi IC50 determination, the drc package of R software was used. Other statistical analyses were performed with student’s T-test or ANOVA depending on the experimental context with GraphPad Prism (RRID:SCR_002798, version 8). (*p<0.05; **p<0.01; ***p<0.001; ****p<0.0001).

## Results

### The BMP4-BMPR1A axis is specifically dysregulated in BLBC and associated with poor prognosis

We previously demonstrated that the BMP2/BMPR1b signaling axis, which is dysregulated in luminal tumors, was directly involved in early steps of MaSC transformation^38^. Given that BMP4 and BMPR1A were also shown to be involved in BLBC, either through their direct effects on BLBC cells or through their dysregulated expression^39,40^, we sought to ascertain their involvement in BLBC emergence. We first compared the transcriptional expression profile of BMPR1A and BMP4 in different breast cancer molecular subtypes and in normal breast tissue (TCGA-Breast cancer and GTEx cohorts). We observed a slight decrease in BMPR1A mRNA expression in all breast cancer subtypes compared to healthy and peritumoral tissue (Figure 1A and Appendix Figure S1A), and a marked decrease in BMP4 mRNA expression specifically in basal-like tumors (Appendix Figure S1B and S1C). Interestingly, the levels of secreted BMP4 protein were similar in BLBC and normal mammary gland, suggesting that tumor cells are not the only source of soluble BMP4 in the tumor microenvironment (Appendix Figure S1D). We then tested whether BMPR1A mRNA levels were correlated with patient prognosis in all breast cancer molecular subtypes and more specifically in basal-like tumors. Interestingly, high BMPR1A expression was associated with a much worse prognosis in the basal-like subtype, but with a slightly better prognosis in all subtypes combined (Figure 1B). Gene ontology pathway analyses revealed that elevated BMPR1A levels were correlated with an increased expression of genes related to chromatin remodeling, migration and response to DNA damage (Figure 1C). Given the link between the pathways identified by GO and DNA repair pathways, including HR, we wondered whether BMPR1A levels were associated with BRCA1 expression. We used BRCA1 wild-type triple-negative breast cancer tumors for which the basal cytokeratin 14 (CK14) expression, highly specific to basal-like breast cancers^43–45^, was determined (Appendix Figure S2A). We then quantified BMPR1A protein expression by immunohistochemistry (Figure 1D) and BRCA1 mRNA expression by RT-qPCR. showed We found an inverse correlation between BRCA1 mRNA and BMPR1A protein levels (Figure 1E). Furthermore, this correlation was conserved after adjustment for age, stage and CK14 membrane expression (Appendix Figure S2B).

**Figure 1.**
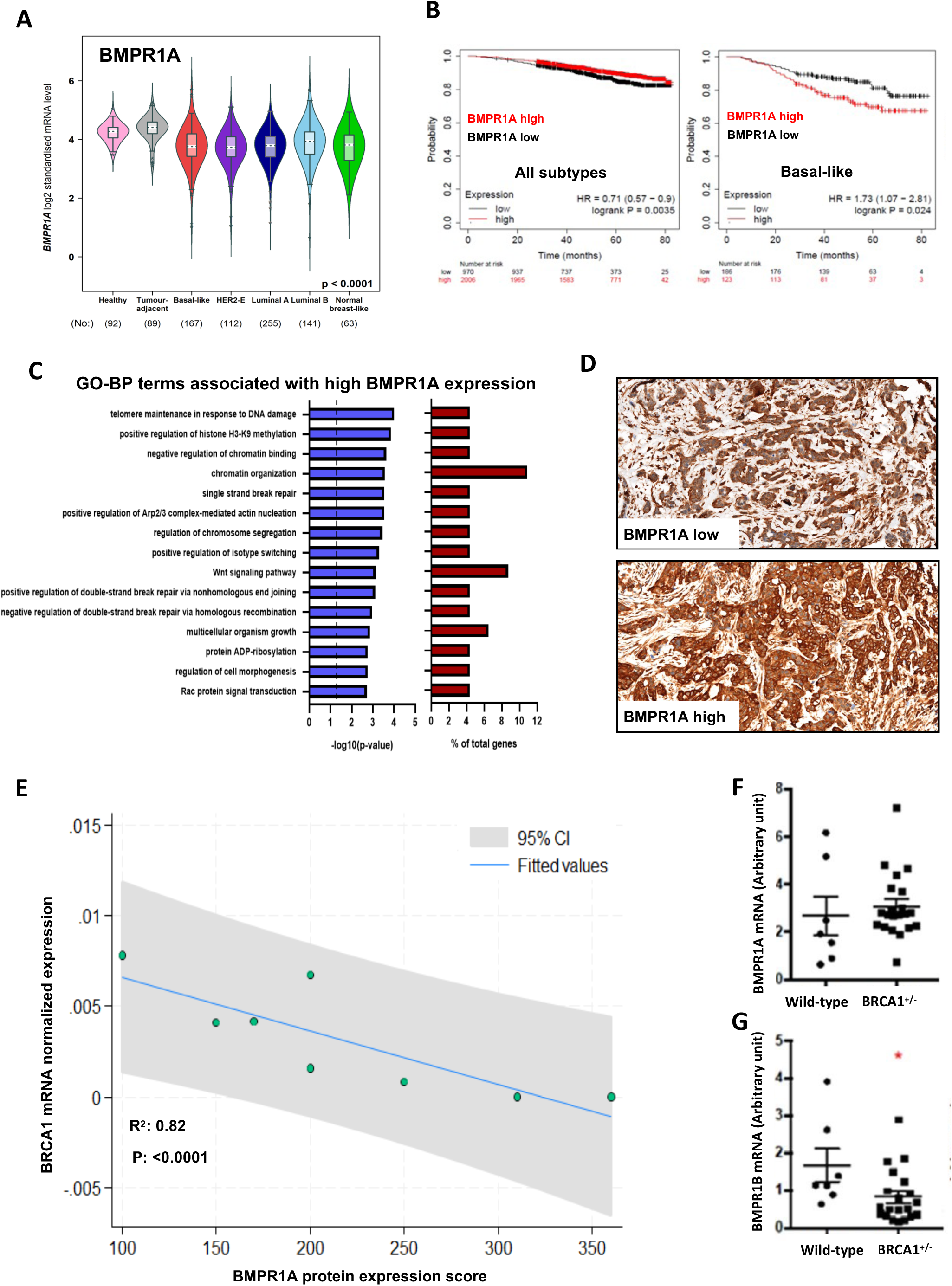
(**A**) BMPR1A mRNA level in breast cancer molecular subtypes, compared with healthy and peri-tumoral tissues in the TCGA-Breast Cancer cohorts. (**B**) Overall survival in function of BMPR1A mRNA level in basal-like cancers (left panel) and all molecular subtypes (right panel) in the TCGA cohort. (**C**) Gene ontology analyses of genes differentially expressed between tumors with high or low BMPR1A expression (TCGA cohort). Statistical significance is plotted on the left and the percentage of differentially expressed genes on the right. (**D**) Representative images of low and strong BMPR1A IHC staining in TNBC. Magnification x20. (**E**) Correlation between BRCA1 mRNA level (normalized on ACTB1 mRNA level) and BMPR1A protein expression (score calculated as the percentage of positive cells x IHC staining intensity). (**F**) RT-qPCR quantification of BMPR1A mRNA level in healthy BRCA1 mutated tissues, compared to healthy wild-type control tissues. Error bars represent standard error to the mean. (**G**) As in **F** for BMPR1B transcriptional expression.

To ascertain at which stage of cell transformation BMP signaling played a role, we compared expression of BMPR1A and BMP4 mRNA in healthy mammary epithelial cells from normal donors and from healthy donors harboring the germinal BRCA1 mutation, and strongly predisposed to developing BLBC^46^. We observed no significant difference in BMPR1A expression (Figure 1F), albeit BRCA1-mutated healthy mammary cells had a lower (yet non-significant) BMP4 expression compared to wild-type samples (Appendix Figure S2C), though this was not confirmed at the secreted protein level (Appendix Figure S2D). In addition, we observed a significant decrease in BMPR1B expression in BRCA1 mutated healthy mammary cells, suggesting that BMPR1A signaling may be dominant over BMPR1B in these cells (Figure 1G). These data suggest that alterations of the BMP pathway are associated with BRCA1 deficiency at a very early, pre-transformation stage. Collectively, our results show that BMPR1A expression is correlated with poor prognosis and is inversely correlated to BRCA1 mRNA level in BLBC.

### BMP4 induces a switch from luminal to myoepithelial differentiation through BRCA1 repression

We next investigated what could be the link between BMP4/BMPR1A signaling and BRCA1 function. We initially conducted an epithelial colony-forming assay (E-CFC) using healthy mammary epithelial cells isolated from patients with BRCA1 haplo-insufficiency (BRCA1), who had undergone prophylactic mastectomy compared to epithelial cells from BRCA1 wild-type (WT) healthy tissue, and observed an increase in basal-like myoepithelial progenitors at the expense of the luminal cells (Appendix Figure S3A, Figure 2A), which is consistent with the previously known role of BRCA1 in luminal differentiation. We then performed functional analyses using MCF10A cells, a human immortalized but not transformed model of mammary stem cells and progenitors^47^. A three-day BMP4 treatment of MCF10A cells induced a similar switch from luminal to myoepithelial progenitors while the total number of colony-forming cells only marginally decreased, suggesting that some luminal progenitors adopt a myoepithelial fate under BMP4 signaling (Figure 2B and Appendix Figure S3B). To determine whether BRCA1 was involved in this change in cell MCF10A cell fate, we developed a doxycycline-inducible shBRCA1 model, which we validated at the transcriptional and protein levels (Figure 2C and 2D). BRCA1 repression in MCF10A cells specifically induced an increase in myoepithelial progenitors at the expense of luminal progenitors, similarly to BMP4 treatment (Figure 2E and Appendix Figure S3C). Interestingly, both BMP4 treatment and BRCA1 knock-down induced a significant increase in the frequency of mammosphere-forming cells, indicative of an increase in immature cells (Appendix Figure S3D and S3E).

**Figure 2.**
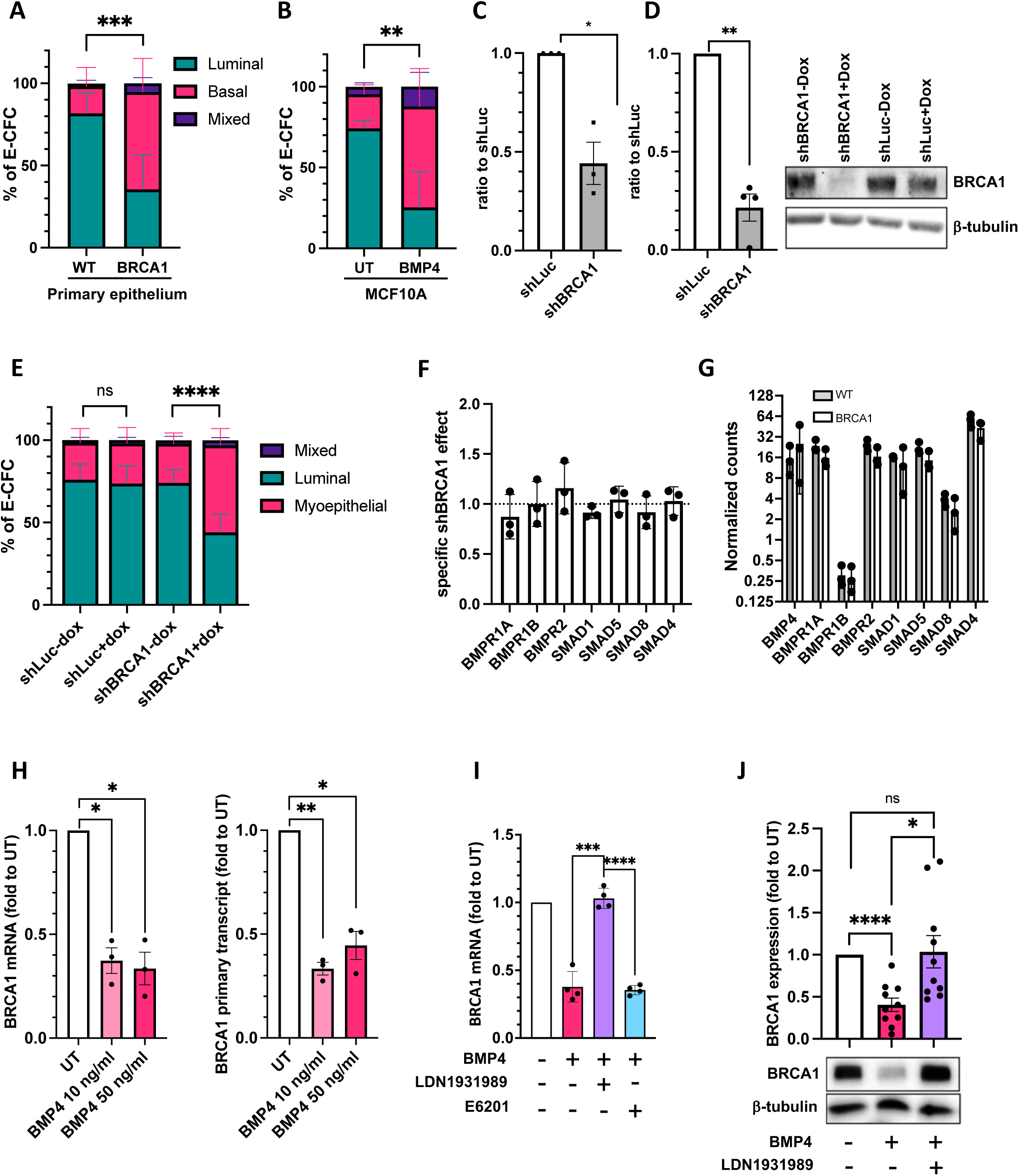
(**A**) Percentage of colonies obtained in E-CFC assays using healthy BRCA1 (BRCA1) mutated or wild-type (WT) mammary epithelial cells. (**B**) Percentage of colonies obtained in E-CFC assays using MCF10A cells exposed or not to BMP4 (10 ng/ml) for 48 hours. (**C**) RT-qPCR quantification of BRCA1 mRNA level in the MCF10A cell line after shBRCA1 induction. Expressed as a ratio between doxycycline effect in shBRCA1 and shLuc cells. (**D**) Quantification and representative image of western-blot analysis of the BRCA1 protein level in the MCF10A cell line after shBRCA1 induction, normalized to the control shLuc (left panel). (**E**) Percentage of colonies obtained in E-CFC assays using MCF10A shBRCA1 or shLuc cells after 48 hours of treatment with or without doxycycline. (**F**) RT-qPCR analyses of the expression of BMP pathway genes following shBRCA1 induction in MCF10A cells. Expressed as a ratio between doxycycline effect in shBRCA1 and shLuc cells (**G**) mRNA levels of genes belonging to the BMP pathway in WT or BRCA1-mutated primary mammary epithelial cells measured by RNA-seq. (**H**) RT-qPCR analysis of BRCA1 mRNA (left panel) and primary transcript (right panel) expression in MCF10A cells exposed for 48 hours to BMP4 (10 and 50 ng/ml). (**I**) RT-qPCR analysis of BRCA1 mRNA expression in response to 48 hours exposure to BMP4 (10 ng/ml) +/- LDN-193189 (100 ng/ml) or E6201 (100nM) added 2 hours before BMP4. (**J**) Quantification and representative picture of western-blot analysis of BRCA1 protein level following a 48 hours treatment with BMP4 (10 ng/ml) +/- LDN-193189 (100 ng/ml) added 2 hours before BMP4. Error bars represent the standard deviation. Statistically significative differences are shown *p<0.05, **p<0.01, ***p<0.001,****p<0.0001.

Given the similar effects observed between BMP4 treatment and BRCA1 knock-down, we wondered if they interacted at a functional level. To test this hypothesis, we measured the specific effects of shBRCA1 on the expression of several genes involved in BMP4 signaling in MCF10A cells by qRT-PCR, including BMP receptors, SMAD1, 5 and 8 effectors, SMAD and the SMAD4 co-SMAD of the BMP and TGF signaling. No significant impact of BRCA1 knock-down was observed on any of these elements at the transcriptional level (Figure 2F), which was further confirmed in RNA-seq data from wild-type of BRCA1-mutated healthy mammary cells regardless of their BRCA1 genomic status (Figure 2G). We then evaluated if BMP4 signaling impacted BRCA1 expression and saw that MCF10A cells that were exposed to BMP4 for 48 h displayed over 50% BRCA1 repression at the mRNA expression (Figure 2H, left panel) and primary transcript (Figure 2H, right panel) levels, suggesting a BMP4-mediated BRCA1 transcriptional repression. In addition, this repression of BRCA1 mRNA expression was reversed using LDN-193189 (a BMPR1A and BMPR1B inhibitor) but not by E6201 (an inhibitor showing a 100-fold higher repression of BMPR1B than of BMPR1A), suggesting the involvement of BMPR1A in BMP4 effect on BRCA1 expression (Figure 2I). Moreover, BRCA1 repression by BMP4 was also observed at the protein level and was completely reversed by LDN-193189 (Figure 2J). Altogether, these results indicate the involvement of BMP4 and BMPR1A in the repression of BRCA1.

### BMP4 induces aberrant RAD51 and 53BP1 recruitment after DNA double-strand breaks

Owing to the well-known involvement of BRCA1 in the most error-free DNA double-strand break (DSB) repair pathway, i.e., homologous recombination (HR), we wondered whether BMP4 affected HR proficiency. Apart from BRCA1, HR requires several other effectors such as RAD51^19^. RAD51 binding on DSB is a reliable marker of HR proficiency^48–52^. Conversely, NHEJ is an error-prone repair mechanism that can be evidenced by 53BP1 recruitment at the DSB^19^. We first validated this in MCF10A cells after 48 hours of shRNA-mediated BRCA1 knock-down before inducing DSB using a 2 Gy irradiation. We measured DNA DSB using γH2AX staining, HR proficiency using RAD51 staining, and NHEJ activity using 53BP1 staining before irradiation and at 4 time-points after irradiation. We saw no significant differences in the number of γH2AX foci at any time-point after irradiation compared to control Luciferase shRNA (Figure 3A), though as expected RAD51 recruitment was markedly decreased at 1 and 4 hours after irradiation (Figure 3B, left panel). We saw no significant differences in 53BP1 recruitment albeit a slight tendency to increase was observed with the BRCA1 shRNA compared to the Luciferase shRNA (Figure 3B, right panel). Similar responses were observed when comparing MCF10A-shBRCA1 with or without doxycycline and no significant effect of the doxycycline was observed in the MCF10A-shLuc cells (Appendix Figures S4 and S5). These results suggest that BRCA1 knock-down in our cell model leads as expected to a HR deficiency likely compensated by the NHEJ pathway. We then performed a similar experiment with MCF10A cells treated or not with BMP4 for 48 hours before irradiation. BMP4 induced no significant differences in the number of γH2AX foci at any time-point after irradiation (Figure 3C and Appendix Figure S6A). Surprisingly, BMP4 treatment led to a significant increase in both RAD51 (Figure 3D, left panel and Appendix Figure S6B) and 53BP1 (Figure 3D, right panel and Appendix Figure S6C) without increasing the total amount of RAD51 protein (Appendix Figure S6D). Taken together, these data support the hypothesis that immature mammary cells have an altered HR in response to BMP4-mediated BRCA1 repression, with an unexpected increase in both RAD51 and 53BP1 recruitment despite the marked decrease in BRCA1 expression.

**Figure 3.**
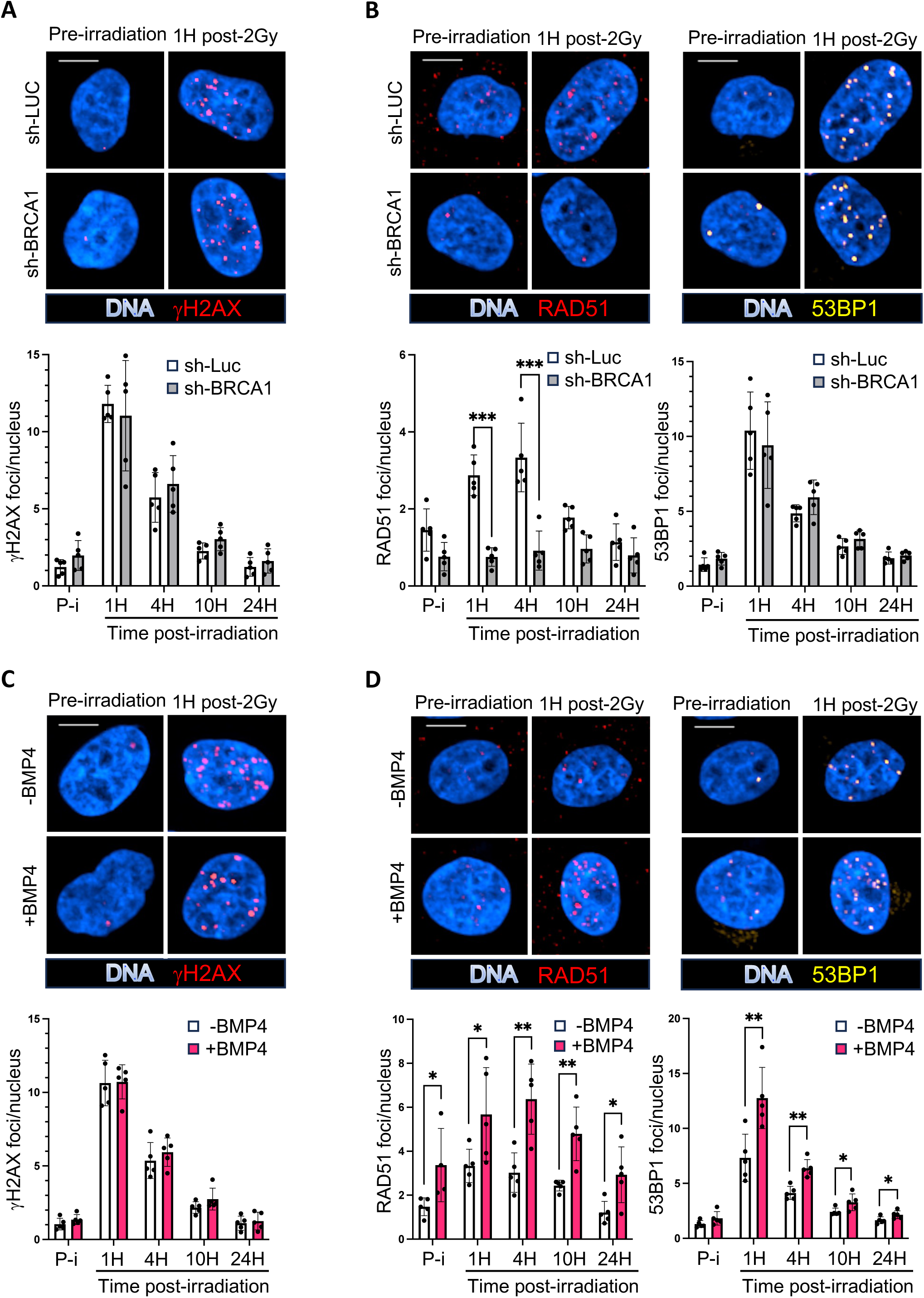
(**A**) Immunofluorescence detection of γH2Ax foci following a 2 Gy irradiation of MCF10A cells after 2 days of doxycycline-induced expression of a shLuc or shBRCA1. The pictures in the upper panel show representative nuclei before irradiation or 1H after irradiation. DNA is stained in blue and γH2Ax foci in red. Pictures were taken with an Opera confocal microscope at 40x magnification and the scale bar represent 10 μm. The lower panel is a quantification of the average number of γH2Ax foci per nucleus in the indicated shRNA expressing cells before (P-i) or after the indicated times after irradiation. (**B**) As in (A) for immunofluorescence detection of RAD51 foci (in red) or 53BP1 foci (in yellow). Both types of foci were detected following a co-staining but each channel is shown independently for clarity. (**C**) Immunofluorescence detection of γH2Ax foci after a 2 Gy irradiation of MCF10A cells following 2 days of culture in presence or absence of BMP4 (10 ng/ml). The pictures in the upper panel show representative nuclei before irradiation or 1H after irradiation. DNA is stained in blue and γH2Ax foci in red. Pictures were taken with an Opera confocal microscope at 40x magnification and the scale bar represent 10 μm. The lower panel is a quantification of the average number of γH2Ax foci per nucleus following the indicated treatment before (P-i) or after the indicated times after irradiation. (**D**) As in (C) for immunofluorescence detection of RAD51 foci (in red) or 53BP1 foci (in yellow). Both types of foci were detected using a co-staining but each channel is shown independently for clarity. Error bars show the standard deviation (n=5). *p<0.05, **p<0.01, ***p<0.001.

### The BMP4-BMPR1A signaling axis induces an increased sensitivity to PARPi

We then explored if the increased RAD51 and 53BP1 recruitment induced by BMP4 to DSBs functionally affected HR. We used inhibitors of PARP enzymes (PARPi) that impair DNA single-strand break repair and result in DSB accumulation. In HR-deficient cells, this leads to cell death by synthetic lethality^53,54^, hence, PARPi are very effective at treating certain cancers with HRD, such as ovarian and breast cancers with BRCA1/2 mutations^55–62^. Sensitivity to PARP inhibitors can therefore be used as a marker of HRD. As a control of our ability to detect HRD through PARPi sensitivity, we first measured the viability of MCF10A cells exposed to increasing doses of PARPi (Olaparib) following the shRNA-mediated knock-down of BRCA1. BRCA1-deficient cells displayed a greater sensitivity to Olaparib compared to cells bearing a control shRNA luciferase lentivector (Figure 4A). We then performed the same assay by exposing MCF10A cells to BMP4, in the presence or not of LDN-193189, and observed a similar increase in sensitivity to Olaparib (Figure 4B and 4C), which was lost following BMPR inhibition. To exclude of-targets effects of Olaparib, we conducted the same experiments with another PARPi, Talazoparib, and obtained the same results (Figure 4D). Next, we tested the involvement of the BMPR1A receptor in this BMP4-mediated effect and generated a BMPR1A-overexpressing MCF10A cell line (MCF10A-R1A+), which was validated at both transcriptional and protein levels compared to MCF10A cells transduced with an empty vector (MCF10A-EV) (Figure 4E and 4F). BMPR1A over-expression, with or without BMP4, induced a similar increase in Olaparib sensitivity that was once again reversed by LDN-193189 (Figure 4G).

**Figure 4.**
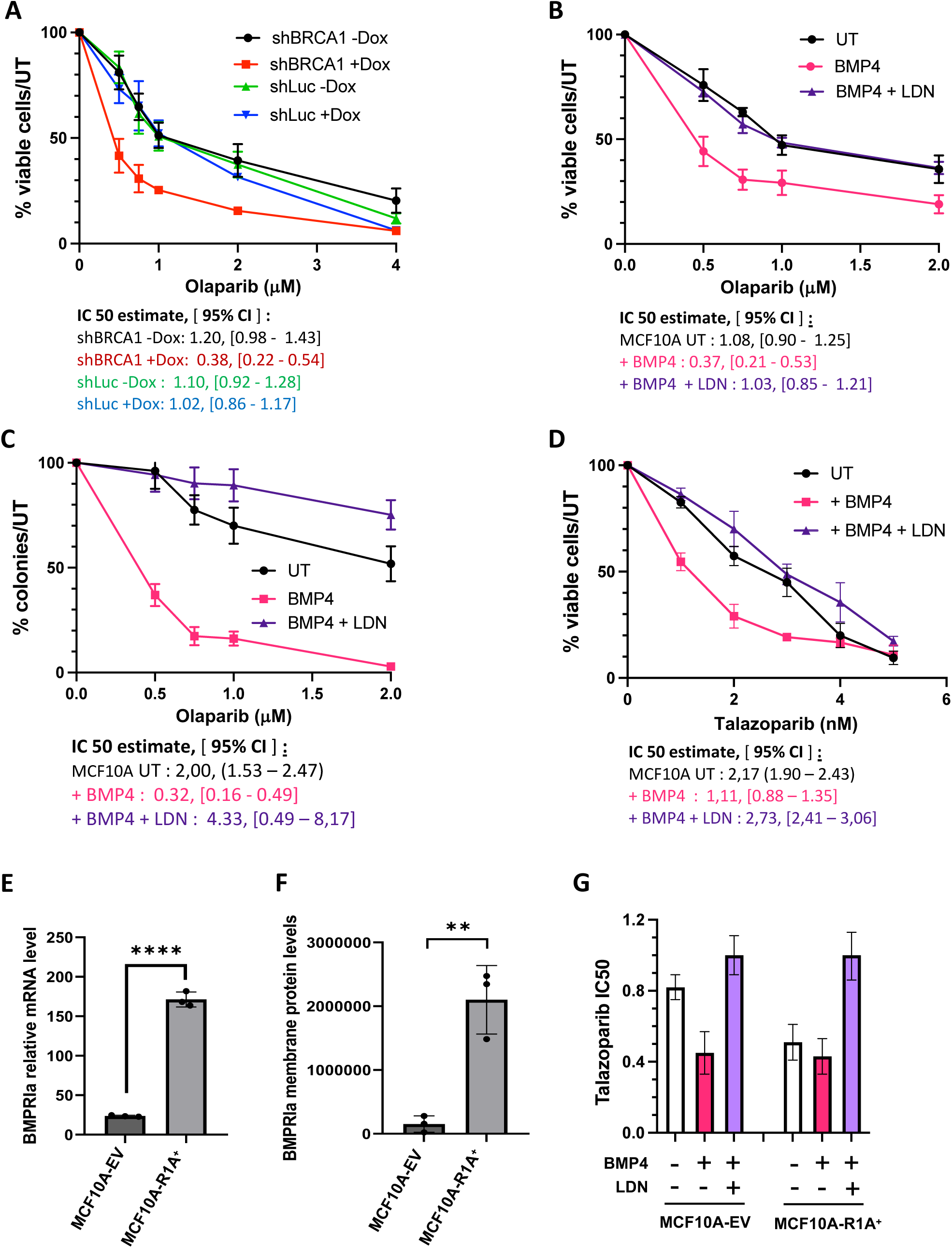
(**A**) Proliferation of MCF10A shBRCA1 or shLuc cells and exposed for 7 days to indicated Olaparib concentrations with or without doxycycline induction. IC50 estimates with confidence intervals are shown below. (**B**) Proliferation assay of MCF10A cells treated or not with BMP4 (10 ng/ml) or BMP4+LDN-193189 (100 ng/ml) and with the indicated doses of Olaparib for 7 days. Representation as in **A**. (**C**) Clonogenic assay of MCF10A cells exposed or not to BMP4 (10ng/ml) or BMP4+LDN-193189 (100 ng/ml) and the indicated doses of Olaparib for 7 days. (**D**) Proliferation assay of MCF10A cells exposed or not to BMP4 (10ng/ml) or BMP4+LDN-193189 (100 ng/ml) and the indicated doses of Talazoparib for 7 days. (**E**) BMPR1A mRNA level in BMPR1A-overexpressing MCF10A (MCF10A-R1A^+^) compared with empty-vector control cells (MCF10A-EV). (**F**) Flow-cytometry analysis of membrane BMPRIA protein expression in BMPR1A-overexpressing MCF10A (MCF10A-R1A^+^) compared with empty-vector control cells (MCF10A-EV). (**G**) Mean IC50 of Talazoparib in MCF10A-R1A^+^ or MCF10A-EV control cells, exposed or not to BMP4 (10ng/ml) and LDN-193189 (100 ng/ml). Error bars represent the 95% confidence interval (n=5).

Altogether these data suggest that BMP4, through its binding to BMPR1A, modulates HR proficiency, likely through BRCA1 repression.

### BMP4 allows the persistence of a fraction of genetically-unstable immature cells at risk of basal-like transformation

Preclinical data for Olaparib in BRCA1-deficient cells previously showed that its antiproliferative effect was due to cell cycle arrest in G2/M and apoptosis^53^. Here, we thus evaluated the level of apoptosis and fate of cells following exposure to BMP4 exposure and treatment with Olaparib. This co-treatment induced a significant increase in apoptosis compared to controls, which was largely lost following the addition of LDN-193189 (Figure 5A). Next, we quantified the effects on the cell cycle, through the incorporation of a Fluorescent and Ubiquination-based Cell Cycle Indicator (FUCCI) system ^63–66^ to generate MCF10A-FUCCI cells. We observed a significant accumulation of cells in G0/early G1 compared to untreated MCF10A cells, suggesting a slowing down of the cell cycle (Figure 5B and Appendix Fig S7A). Interestingly, BMP4 or Olaparib alone also resulted in G0/early G1 arrest but to a lesser extent. This accumulation of cells in G0/G1 could be related to an immature phenotype. To test this hypothesis, we measured by mammosphere and E-CFC assays the stem cell/progenitor ratio in each experimental condition, and obtained a significantly higher ratio (over 6-folds) in surviving cells co-treated with BMP4 and Olaparib compared to untreated cells or cells treated with BMP4 or Olaparib alone (Figure 5C). Hence, this co-treatment led to the generation of cells with a stronger stemness phenotype, which was reversed by the addition of LDN-193189 (Figure 5C).

**Figure 5.**
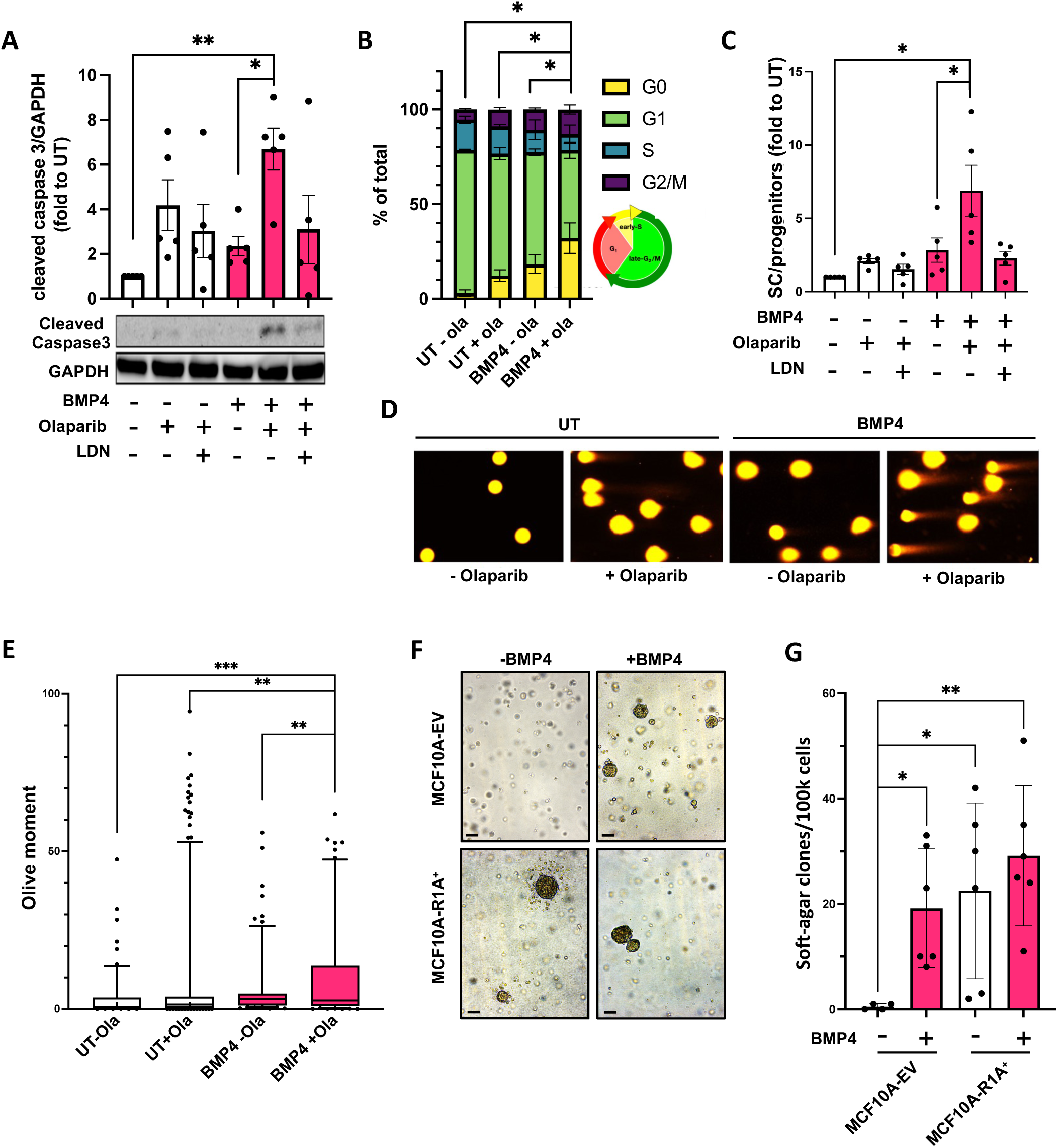
(**A**) Western-blot quantification and representative picture of cleaved caspase-3 protein level in MCF10A cells treated or not with BMP4 (10 ng/ml), LDN-193189 (100 ng/ml) and Olaparib (0.75 μM) for 7 days. GAPDH is used as a loading control. (**B**) Distribution in the different phases of the cell cycle of MCF10A-Fucci cells treated or not with BMP4 (10 ng/ml) and with Olaparib (0.75 μM) for 7 days. (**C**) MCF10A cells were treated or not with BMP4 (10 ng/ml), LDN-193189 (100 ng/ml) or Olaparib (0.75 μM) for 7 days and the ratio between mammosphere-forming cells and colony-forming cells frequencies determined. (**D**) Representative pictures of a neutral comet assay on MCF10A cells exposed or not to BMP4 (10ng/ml) or/and Olaparib (0.75 μM) after 7 days of treatment. (**E**) Quantification of the olive moment in the Comet assay shown in (D). (**F**) Representative pictures of colonies (magnification: 10x) and quantification (**G**) of the number of colonies formed in a soft-agar culture by MCF10A cells overexpressing or not BMPR1A and exposed or not to BMP4 for 20 weeks. Error bars represent standard error to mean. *p<0.05, **p<0.01, ***p<0.001.

The persistence of such stem-like cells with reduced BRCA1 expression levels and altered HR could lead to a genetically unstable phenotype. We evaluated the level of DNA damage in persistent cells after BMP4 and Olaparib treatment using a neutral comet assay to detect unresolved DNA double-strand breaks, as well as an alkaline comet assay to include single-strand breaks. In persistent cells, co-exposure to BMP4 and Olaparib induced a higher number of DSBs (Figure 5D, 5E and Appendix Figure S7B and S7C). Finally, to verify whether long-term BMP4-BMPR1A signaling, independently of any exogenous inducer of DNA damage, could induce transformation, we exposed BMPR1A-overexpressing MCF10A cells and their control to BMP4 in an 18-week culture and performed soft agar assays. BMP4 exposure or BMPR1A overexpression generated a greater number of soft-agar colonies than in the control condition (Figure 5F, 5G), demonstrating that sustained BMP4/BMPR1A signaling in immature mammary cells can lead to their transformation, possibly associated with an increased genomic instability caused by BRCA1 repression.

## Discussion

Basal-like breast cancers present significant clinical challenges due to their aggressive nature and limited targeted therapeutic options. A substantial proportion of these cancers are not associated with BRCA1 mutations, complicating treatment strategies and underscoring the need for a better understanding of their biology. Poly (ADP-ribose) polymerase inhibitors (PARPi) have emerged as a promising therapeutic option, particularly in tumors with deficiencies in homologous recombination repair, offering a synthetic lethality approach to target these malignancies ^67^. However, the persistence of cancer stem cells raises critical challenges to achieve durable remission, as these cells are often resistant to therapies, including PARPi, and contribute to disease recurrence and metastasis^67^. In this context, our study provides new insight into early mechanisms that could promote basal-like carcinogenesis in the absence of BRCA1 mutations. We reveal the role of the mammary gland microenvironment, notably the BMP4-BMPR1A axis, as high BMPR1A expression at the mRNA level was correlated with poor patient prognosis specifically in BLBC. Moreover, we show that BMPR1A protein expression is inversely correlated to BRCA1 mRNA expression and indicative of bad prognosis.

We studied the effects of the BMP4/BMPR1A axis using the MCF10A cell line, which is an immortalized but not transformed model (CDKN2A homozygous deletion, cMYC amplification)^68^. This cellular model also represents a heterogeneous population of human mammary stem-like cells able to generate bipotent or unipotent luminal and basal progenitors in response to microenvironmental cues^38^. Here, we saw that BMP4 induced a transcriptional repression of BRCA1, associated with and sufficient to induce a switch towards basal progenitor differentiation. Accordingly, we evidenced a role for BRCA1 repression on progenitor fate choice in the mammary gland toward an enrichment in basal differentiation and stemness, reminiscent of the impact of BMP4 on human primary mammary stem cells^38,69,70^. This is also consistent with several previous studies carried out *ex vivo* under BRCA1 haploinsufficiency^71–73^ and *in vitro*^74,75^ that have all shown a role for BRCA1 in differentiating breast luminal progenitors. Therefore, for the first time, our data link basal differentiation mediated by BRCA1 repression and BMP signaling in non-transformed cells.

Poly (ADP-ribose) polymerase (PARP) 1 and PARP2 are enzymes involved in the response to DNA damage, in particular DNA single-strand break repair by base excision. PARP inhibitors (PARPi) are competitive inhibitors of PARP1 and PARP2. This results in the non-resolution of single-strand DNA breaks and the trapping of PARP enzymes on damaged DNA. The resulting blockage of the replication fork when the cell enters the replicative phase leads to the formation of DSBs. In the context of a homologous recombination deficiency (HRD), this damage of the replication fork cannot be reliably repaired. This leads to the activation of other error-prone repair systems favoring the accumulation of genetic lesions leading to cell death by synthetic lethality^53,54^. PARP inhibitors (PARPi) are very effective in several tumors with know HRD, notably certain epithelial ovarian cancers and certain breast cancers with a constitutional BRCA1/2 deleterious mutation^55–62^. Here, we demonstrated that the BRCA1 repression induced by BMP4 was sufficient to induce HRD as evidenced by an increased sensitivity to PARPi. Moreover, while BMP4-induced HRD and Olaparib-induced DNA damage have an additive effect on apoptosis-induced cell death, our data also revealed an accumulation of surviving cells in a G0/G1 stage correlated with an increase in immature features. This preferential survival of immature resting cells could constitute a cellular reservoir of preneoplastic cells. Indeed, comet assays further suggest an associated genomic instability due to BMP4-induced HRD and indicated by accumulation of unresolved DSB after PARPi treatment. The notion that only a partial decrease in BRCA1 function can induce a genomic instability that could lead to transformation is supported by other studies. For example, the introduction of a heterozygous BRCA1 mutation (185delAG) into the MCF10A cell line was reported to induce copy number alterations^76^. In addition, healthy BRCA1 mutated tissues have an intermediate level of chromosomal aberrations, between non-mutated healthy control tissues and BRCA1 mutated tumors^77^. Finally, unlike BRCA-mutated ovarian tumors, loss of the BRCA1 WT allele is not necessary for the carcinogenesis of BRCA1-mutated breast tumors^78^. Partial defect in BRCA1 function is thus sufficient to induce a genetic instability before the transformation stage, as well as during transformation. The hyper-recombination phenotype linked to BRCA1 haploinsufficiency could be causing this initial genetic instability^79^. Our data are thus compatible with this hypothesis since we observed that RAD51 was over-recruited to DSBs following BMP4 exposure. It has been proposed that such RAD51 over-recruitment at DSBs could promote non-homologous hyper-recombination and chromosomal aberrations causing loss of heterozygosity and aneuploidy that could participate to carcinogenesis^80–87^. In addition, RAD51 can generate genetic instability caused by non-homologous (or micro-homologous) recombination, a process countered by 53BP1^88^.

In this context, we demonstrate here that chronic exposure to BMP4 could induce transformation. Indeed, long-term treatment of the MCF10A cell line with BMP4 led to their transformation, likely through BMP4-induced BRCA1 repression. This uncovers a new mechanism to establish a BRCAness phenotype in non-transformed cells through a BMP4/BMPR1A-mediated BRCA1 transcriptional repression.

In a physiological context, BMP4 is an important cytokine of the mammary gland microenvironment that induces the basal differentiation of the mammary progenitors^38^, acting at least in part through BRCA1 repression. BMP4 signaling is likely integrated with other cytokines and hormones to ensure the homeostasis and plasticity of the mammary gland during life. BRCAness induced by germline BRCA1 mutations, albeit present in all cells of the body, predispose almost exclusively to cancers of the breast and ovary, two organs where female hormones play a key role. Among these hormones, estrogens and especially estradiol (E2) seems to be of particular interest in this context. Indeed, E2 is involved in luminal differentiation and activates BRCA1 expression, suggesting that BMP4 and E2 may control the balance between basal and luminal differentiation. Any alteration of this balance in favor of BMP4 signaling could lead to increased basal differentiation associated with BRCA1 deficiency and genomic instability. Chronic exposure to environmental pollutants could be an example of perturbation since we previously showed that exposures to bisphenol A or Benzo(A)pyrene, both associated to increased BC risk^89,90^, strongly increase BMPR1A expression and affect stem cells features^91^. This study contributes to a better understanding of the complex regulatory mechanisms controlling mammary cell fate and BRCA1 expression.

In the clinic, PARP inhibitors are used in subsets of BLBC patients with BRCA1 germline mutations but this could represent only a subset of BLBC patients that could benefit from therapies exploiting HRD. Identifying BLBC patients with ongoing HRD due to increased BMP4/BMPR1A signaling and subsequent BRCA1 repression could led to the extension of PARP inhibitors to other patients. In addition, targeting the BMP4/BMPR1A pathway in these BLBC could be an alternative therapeutic option that should be further explored to treat patients with the most aggressive breast cancer subtype.

## Ethic statement

The use of patient samples was validated by the ethical committee of the Centre Léon Bérard under treatment number R201-004-531. All patients previously gave their informed consent for the use of the samples.

## Authors’ Contributions

Simon AHO: Conceptualization, investigation, methodology, formal analysis, writing original draft. Marie PERBET : Investigation. Isabelle TREILLEUX : Formal analysis, investigation. Adrien BUISSON : Investigation. Sandrine JEANPIERRE : Investigation, methodology, Emmanuel DELAY : Conceptualization, supervision, resources. Véronique MAGUER-SATTA : Conceptualization, formal analysis, supervision, funding acquisition, investigation, methodology, writing original draft, project administration. Boris GUYOT: Conceptualization, formal analysis, supervision, funding acquisition, investigation, methodology, writing original draft.

## Acknowledgments

We thank Dr Patrick Mehlen for its support and helpful advices. We thank P. Battiston-Montagne and C. Vanbelle, CRCL-PIC cytometry and imaging platform. We thank B. Manship for proofreading. This study was funded by Convergence PLAsCAN ANR-17-CONV-0002); La Ligue Nationale Contre le Cancer Ain, Rhône and Saône-et-Loire (VMS); Fondation ARC (SFI20111203500, PJA20171206331); the ERiCAN program of Fondation MSD-Avenir (DS-2018-0015) and Associations [Déchaîne Ton Cœur; RUBAN ROSE Prix Avenir 2021; Comité féminin pour le dépistage du cancer du sein 74; and Fondation pour la Recherche Medicale (equipe FRM EQU202203014695).

