## Supplemental Figures S1 to S7 for "The BMP4-BMPR1A axis represses BRCA1 in mammary stem cells and contributes to tumor initiation"

**A**

| BMPR1A | Dunnett-Tukey-Kramer's test: | | | p-value |
|---|---|---|---|---|
| gene-expression comparisons | | | | |
| Basal-like | < | Healthy | ✓ | < 0.0001 |
| Basal-like | < | Tumour-adjacent | ✓ | < 0.0001 |
| Basal-like | = | HER2-E | ✗ | > 0.10 |
| Basal-like | = | Luminal A | ✗ | > 0.10 |
| Basal-like | = | Luminal B | ✗ | > 0.10 |
| Basal-like | = | Normal breast-like | ✗ | > 0.10 |

**B**

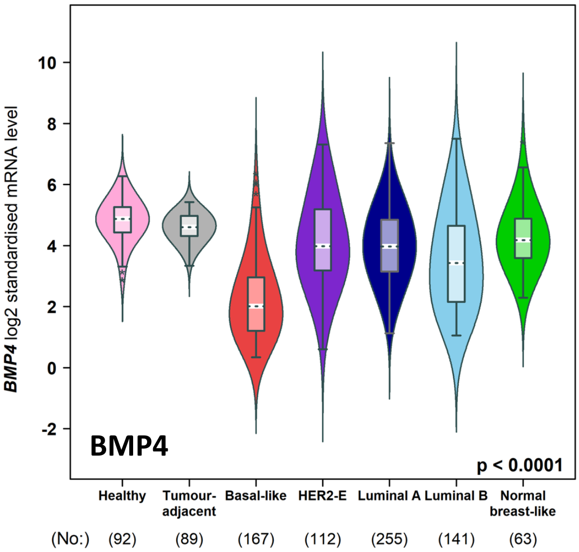


**C**

| BMP4 | Dunnett-Tukey-Kramer's test: | | | p-value |
|---|---|---|---|---|
| gene-expression comparisons | | | | |
| Basal-like | < | Healthy | ✓ | < 0.0001 |
| Basal-like | < | Tumour-adjacent | ✓ | < 0.0001 |
| Basal-like | < | HER2-E | ✓ | < 0.0001 |
| Basal-like | < | Luminal A | ✓ | < 0.0001 |
| Basal-like | < | Luminal B | ✓ | < 0.0001 |
| Basal-like | < | Normal breast-like | ✓ | < 0.0001 |

**D**

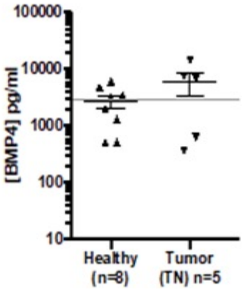

Appendix Figure S2

**A**

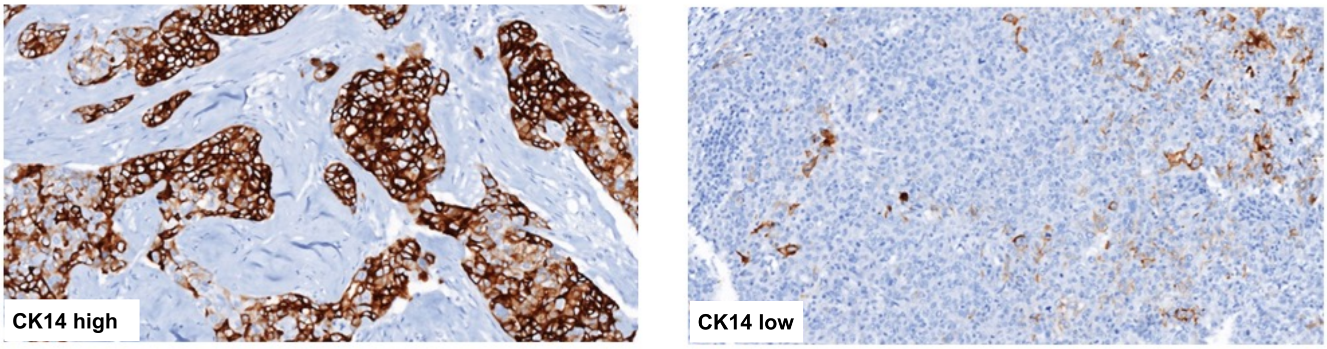


**B**

| Adjustement variable | BMPR1A total R-squared | Coefficient | 95% CI | P-value |
|---|---|---|---|---|
| Age | 0.7148 | -.0000309 | -.0000489<br>-.000013 | 0.007 |
| Stage | 0.6956 | -.0000296 | -.000046<br>-.0000132 | 0.006 |
| CK14 membrane expression | 0.8193 | -.000031 | -.0000384<br>-.0000237 | <0.0001 |

**C**

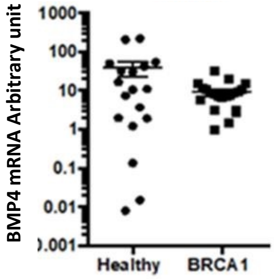


**D**

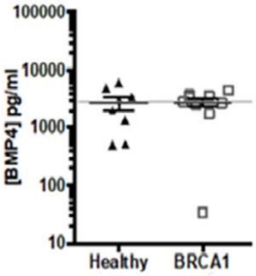

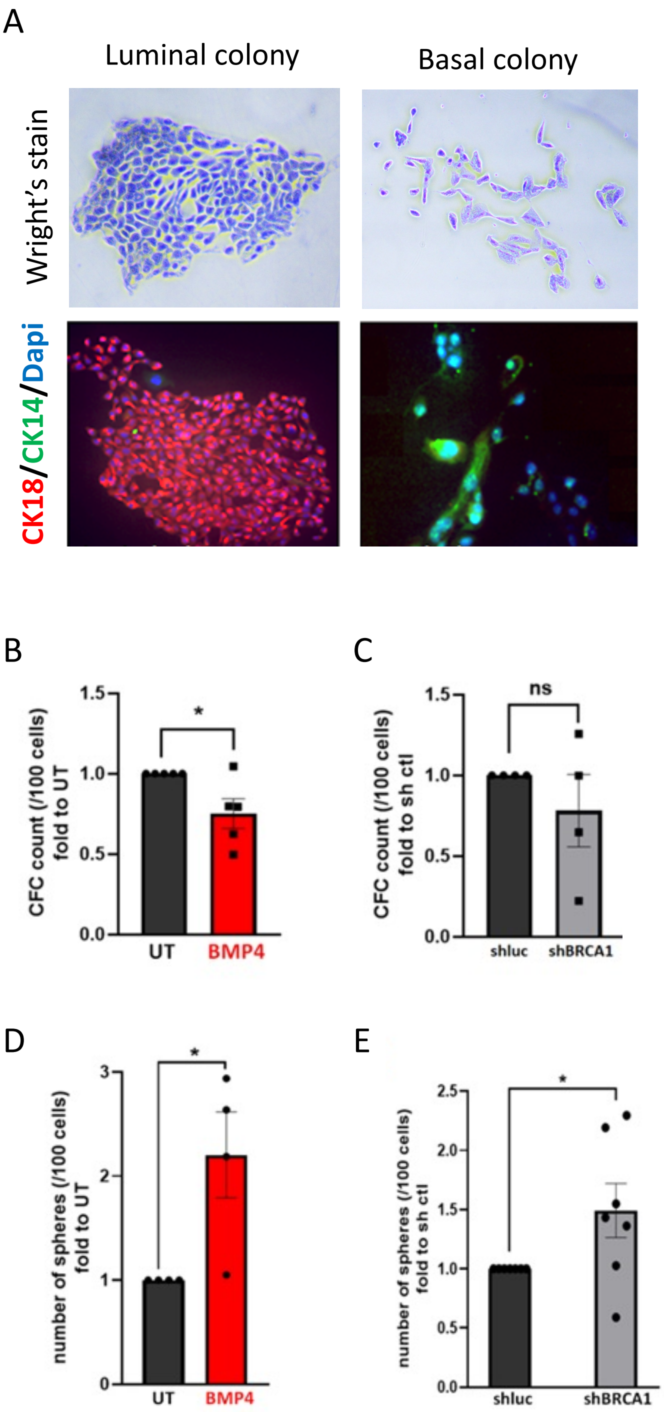
A
Luminal colony
Basal colony
Wright's stain
CK18/CK14/Dapi
B
CFC count (/100 cells) fold to UT
1.5
1.0
0.5
0.0
*
UT
BMP4
C
CFC count (/100 cells) fold to sh ctl
1.5
1.0
0.5
0.0
ns
shluc
shBRCA1
D
number of spheres (/100 cells) fold to UT
3
2
1
0
*
UT
BMP4
E
number of spheres (/100 cells) fold to sh ctl
2.5
2.0
1.5
1.0
0.5
0.0
*
shluc
shBRCA1

Appendix Figure S4

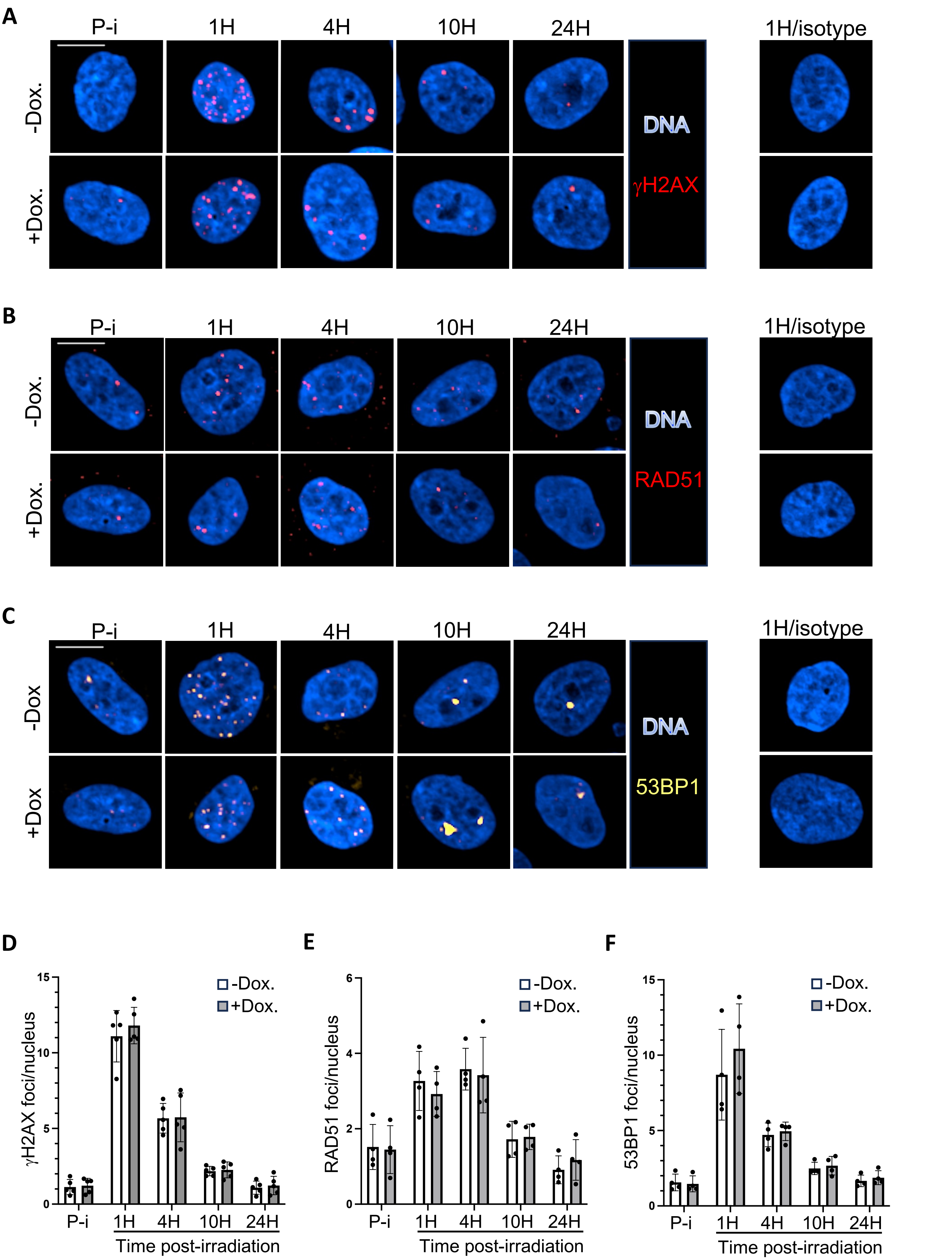

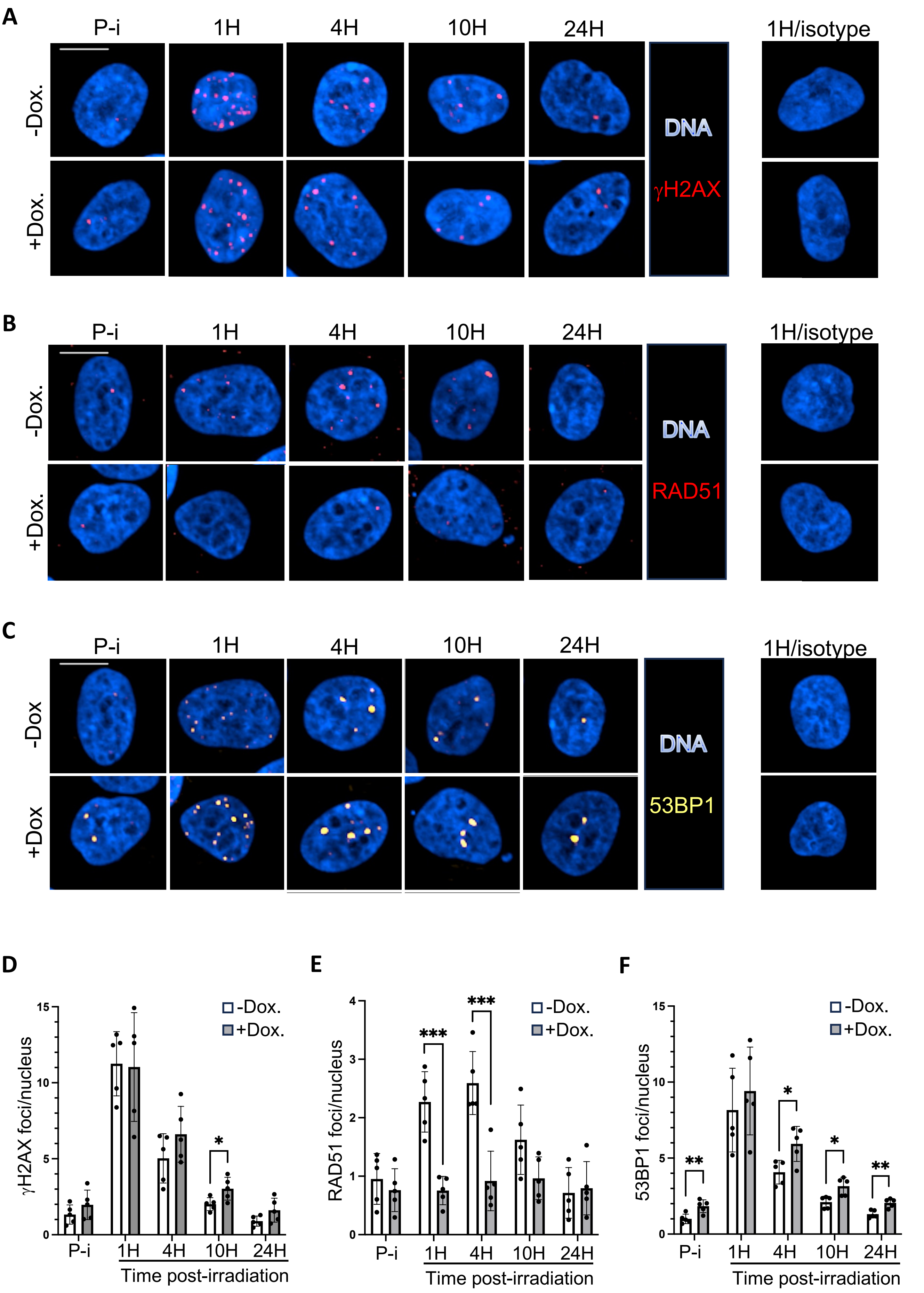
A
P-i
1H
4H
10H
24H
1H/isotype
-Dox.
+Dox.
DNA
γH2AX
B
P-i
1H
4H
10H
24H
1H/isotype
-Dox.
+Dox.
DNA
RAD51
C
P-i
1H
4H
10H
24H
1H/isotype
-Dox
+Dox
DNA
53BP1
D
γH2AX foci/nucleus
-Dox.
+Dox.
P-i
1H
4H
10H
24H
Time post-irradiation
E
RAD51 foci/nucleus
***
***
-Dox.
+Dox.
P-i
1H
4H
10H
24H
Time post-irradiation
F
53BP1 foci/nucleus
**
*
*
**
-Dox.
+Dox.
P-i
1H
4H
10H
24H
Time post-irradiation

Appendix Figure S6

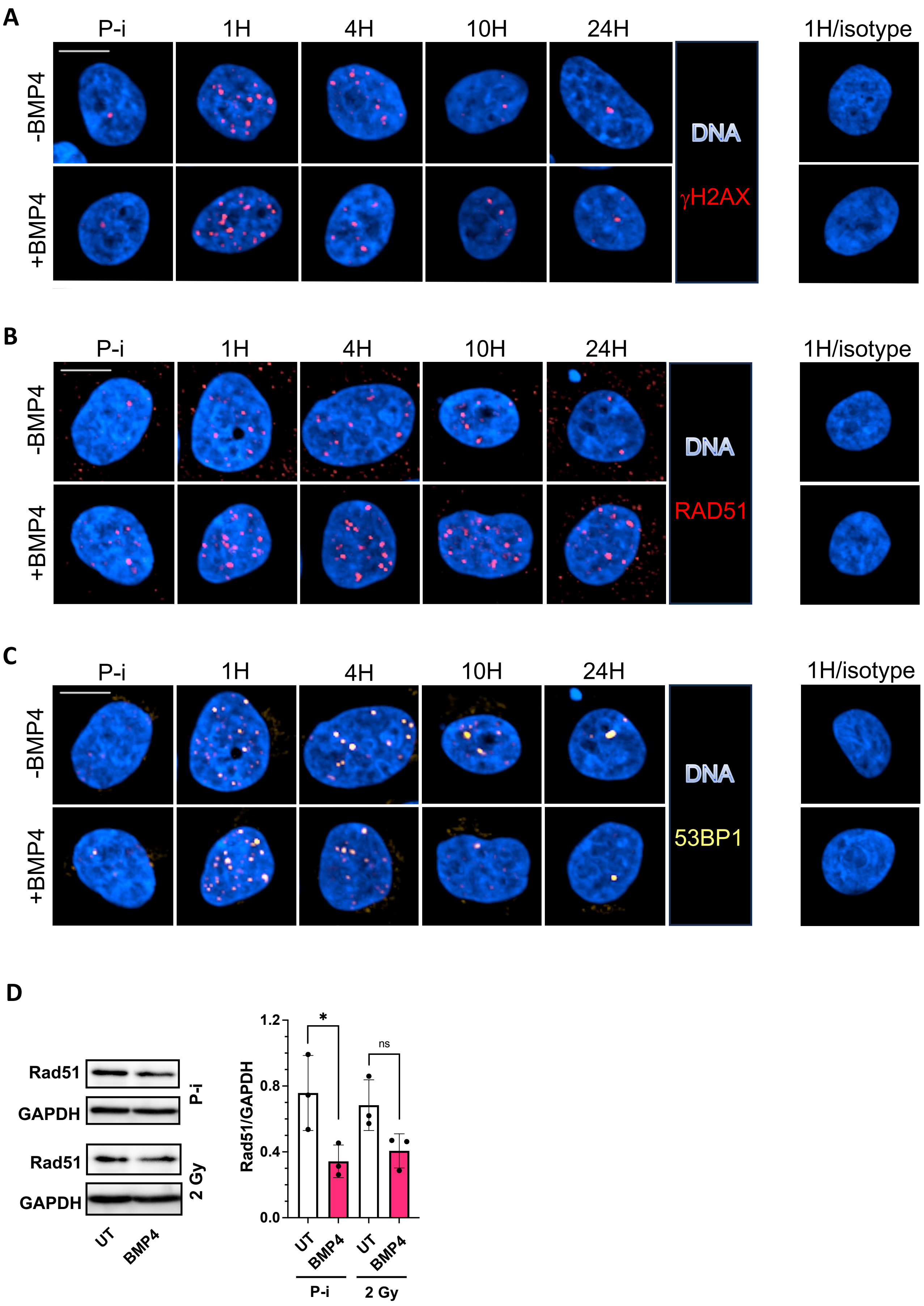

Appendix Figure S7

**A**

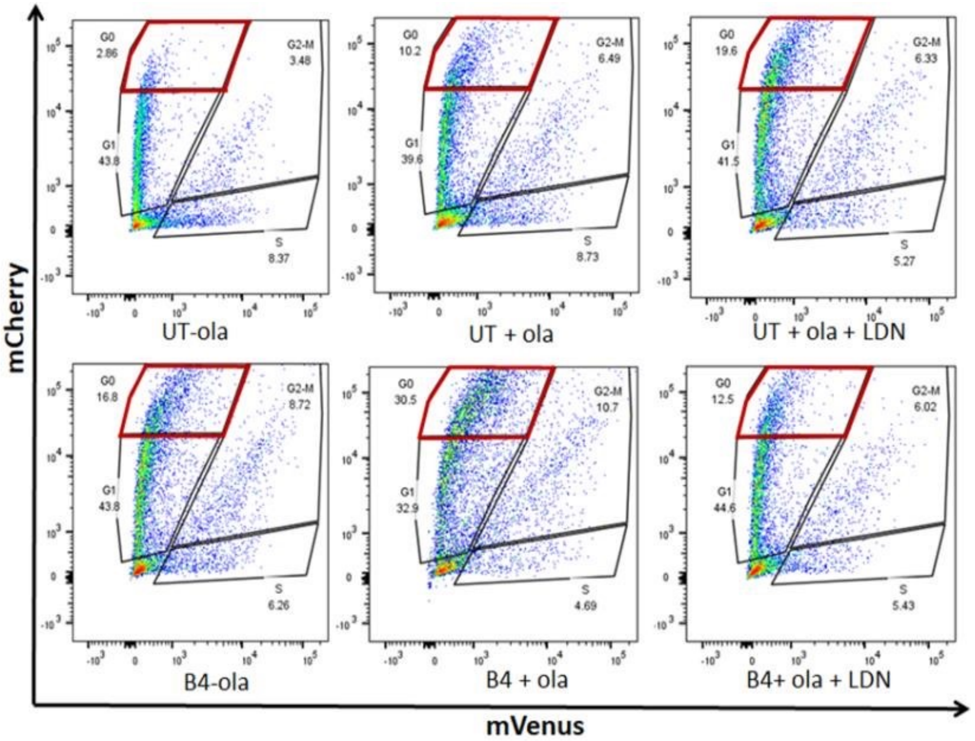


**B**

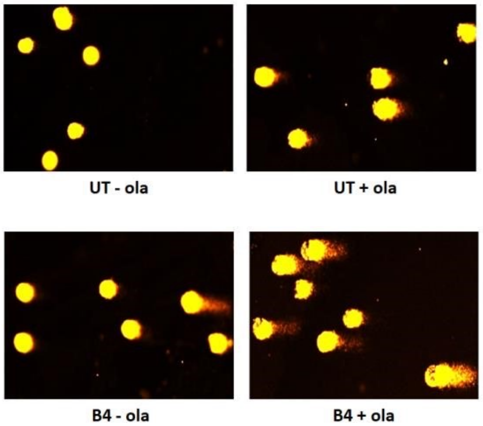


**C**

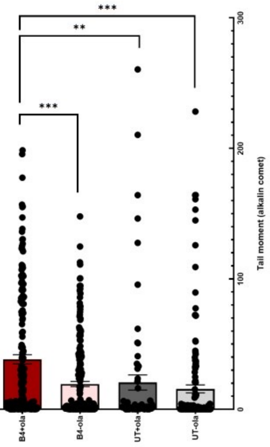

### Appendix figures legends

**Appendix Figure S1. (A)** Dunnet-Tukey-Kramer statistical test of the significativity of the BMPR1A mRNA expression differences between BLBL and the indicated PAM50 breast cancer subtypes in the TCGA cohort. **(B)** BMP4 mRNA expression in different breast cancer molecular subtypes, compared with healthy and peri-tumoral control tissues in the TCGA-Breast Cancer cohort. **(C)** Dunnet-Tukey-Kramer statistical test of the significativity of the BMP4 mRNA expression differences between BLBC and the indicated PAM50 breast cancer subtypes in the TCGA cohort. **(D)** ELISA quantification of soluble BMP4 in healthy mammary tissue and TNBC samples.

**Appendix Figure S2. (A)** Representative high (left picture) and low (right picture) IHC staining of CK14 expression in BRCA1 wild-type TNBC. **(B)** Results of bivariate linear regression models of the correlation between BMPR1A protein and BRCA1 mRNA expression after adjustment for age, stage or CK14 membrane expression. **(C)** RT-qPCR analysis of BMP4 transcriptional expression in healthy BRCA1 mutated tissues, compared to healthy wild-type control tissues. **(D)** ELISA quantification of soluble BMP4 in healthy BRCA1 mutated tissues compared to healthy wild-type control tissues.

**Appendix Figure S3. (A)** Representative Wright's staining (upper row) or immunofluorescence (lower row) of CK18 (red), CK14 (green) and DNA (blue) in the indicated type of colonies obtained following an E-CFC assay in MCF10A. **(B)** Impact of a 72 hours BMP4 exposure of MCF10A on the total number of E-CFC compared to untreated cells. **(C)** Impact of a 72 hours shRNA-mediated BRCA1 knock-down in MCF10A on the total number of E-CFC compared to shLuc control cells. **(D)** Impact of a 72 hours BMP4 exposure of MCF10A on the number of mammosphere-forming cells compared to untreated cells. **(E)** Impact of a 72 hours shRNA-mediated BRCA1 knock-down in MCF10A on the number of mammosphere-forming cells compared to shLuc control cells.

**Appendix Figure S4. (A)** Representative nuclei from an immunofluorescence detection of γH2Ax foci at the indicated time points before (P-i) or following a 2 Gy irradiation of MCF10A cells bearing a doxycycline-inducible shRNA directed against the Luciferase mRNA after 2 days in absence (-Dox) or presence (+Dox) of 400 ng/ml of doxycycline. DNA is stained in blue and γH2Ax in red. Representative nuclei 1 H post-irradiation with the anti-γH2AX antibody replaced by an isotype control are shown on the right panel. Pictures were taken with an Opera confocal microscope at 40x magnification and the scale bar represent 10 μm. **(B)**. As in (A) using an antibody directed against RAD51 (in red). **(C)**. As in (A) using an antibody directed against 53BP1 (in yellow). **(D)**. Quantification of the average number of γH2Ax foci per nucleus in the experiments illustrated in (A). Error bars show the standard deviation (n=5). *$p<0.05$, **$p<0.01$, ***$p<0.001$. **(E)**. As in (D) for the quantification of RAD51 foci illustrated in (B). **(F)**. As in (D) for quantification of 53BP1 foci illustrated in (C).

**Appendix Figure S5. (A)** Representative nuclei from an immunofluorescence detection of γH2Ax foci at the indicated time points before (P-i) or following a 2 Gy irradiation of MCF10A cells bearing a doxycycline-inducible shRNA directed against the *BRCA1* mRNA after 2 days in absence (-Dox) or presence (+Dox) of 400 ng/ml of doxycycline. DNA is stained in blue and γH2Ax in red. Representative nuclei 1 H post-irradiation with the anti-γH2AX antibody replaced by an isotype control are shown on the right panel. Pictures were taken with an Opera confocal microscope at 40x magnification and the scale bar represent 10 μm. **(B)** As in (A) using an antibody directed against RAD51 (in red). **(C)**. As in (A) using an antibody directed against 53BP1 (in yellow). **(D)** Quantification of the average number of γH2Ax foci per nucleus in the experiments illustrated in (A. Error bars show the standard deviation (n=5). *$p<0.05$, **$p<0.01$, ***$p<0.001$. **(E)** As in (D) for the quantification of RAD51 foci illustrated in (B). **(F)** As in (D) for quantification of 53BP1 foci illustrated in (C).

**Appendix Figure S6. (A)** Representative nuclei from an immunofluorescence detection of γH2Ax foci at the indicated time points before (P-i) or following a 2 Gy irradiation of MCF10A cells after 2 days in absence (-BMP4) or presence (+BMP4) of 10 ng/ml of BMP4. DNA is stained in blue and γH2Ax in red.

Representative nuclei 1 H post-irradiation with the anti-γH2AX antibody replaced by an isotype control are shown on the right panel. Pictures were taken with an Opera confocal microscope at 40x magnification and the scale bar represent 10 µm. **(B)** As in (A) using an antibody directed against RAD51 (in red). **(C)** As in (A) using an antibody directed against 53BP1 (in yellow). **(D)** Left panel: representative pictures of western-blot detection of RAD51 before (P-i) or 1 hour following a 2 Gy irradiation of MCF10A cells after 2 days in absence (UT) or presence (BMP4) of 10 ng/ml of BMP4. GAPDH detection is used as a loading control. Right panel: Quantification of the RAD51/GAPDH ratio in the indicated conditions. Error bars show the standard deviation (n=5). *$p<0.05$.

**Appendix Figure S7. (A)** Representative FACS plots of MCF10A-Fucci cells exposed or not to BMP4 (10ng/ml) and LDN-193189 (100 ng/ml) and treated or not with Olaparib (0.75 µM) for 7 days. **(B)** Representative pictures of a basic comet assay on MCF10A cells exposed or not to BMP4 (10ng/ml) or/and Olaparib (0.75 µM) after 7 days of treatment. **(C)** Quantification of the olive moment in the Comet assay shown in (B).
